# Spray-induced gene silencing as a potential tool to control potato late blight disease

**DOI:** 10.1101/2021.02.07.430140

**Authors:** Pruthvi B. Kalyandurg, Poorva Sundararajan, Mukesh Dubey, Farideh Ghadamgahi, Muhammad Awais Zahid, Stephen C. Whisson, Ramesh R. Vetukuri

## Abstract

*Phytophthora infestans* causes late blight disease on potato and tomato and is currently controlled by resistant cultivars or intensive fungicide spraying. Here, we investigated an alternative means for late blight control by spraying potato leaves with double-stranded RNAs (dsRNA) that target *P. infestans* genes that are essential for infection. Through confocal microscopy, we show that the sporangia of *P. infestans* expressing Green Fluorescent Protein (GFP) can take up *in vitro* synthesized dsRNAs homologous to GFP directly from their surroundings, including leaves, which leads to the reduced relative expression of *GFP*. We further demonstrate the potential of spray induced gene silencing (SIGS) in controlling potato late blight disease by targeting developmentally important genes in *P*.*infestans* such as guanine-nucleotide binding (G) protein β-subunit (*PiGPB1*), haustorial membrane protein (*PiHmp1*), cutinase (*PiCut3*), and endo-1,3(4)-β-glucanase (*PiEndo3*). Our results demonstrate that SIGS can be potentially used to mitigate potato late blight; however, the degree of disease control is dependent on the selection of the target genes.

---

Despite causing devastating late blight disease on tomato and potato worldwide, there are few alternatives to plant resistance or chemical control for the plant pathogenic oomycete, *Phytophthora infestans* (Kamoun et al., 2015). Given that *P. infestans* is a fast-growing, highly adaptable filamentous pathogen, traditional breeding for resistance has not proved durable in the field (Leesutthiphonchai et al., 2018; Whisson et al., 2016). One of the most effective control methods available for late blight control is intensive fungicide spraying, costing billions of dollars to potato and tomato growers annually, and raises serious environmental concerns. Moreover, *P. infestans* has overcome resistance to some of the fungicides in use (Schepers et al., 2018). Many countries have increased the stringency of regulations governing approval of agrochemicals, potentially limiting the selection of effective fungicides available. There is thus an urgent need to develop alternative means for pathogen control.

RNA interference (RNAi) is a conserved cellular defence mechanism mediated by double-stranded RNA (dsRNA) regulating protein expression through targeted destruction or modulation of mRNAs (Ghildiyal & Zamore, 2009; Hannon, 2002; Huang et al., 2019; Malone & Hannon, 2009), or modification of chromatin (Van Wolfswinkel & Ketting, 2010). While RNAi is a fundamental cellular defence mechanism against invading pathogens, introducing *in vitro* synthesized dsRNAs or producing the molecules *in planta* exploits this natural cellular reaction as a crop management strategy (Huang et al., 2019) and is a promising new method for controlling plant diseases.

Plant transgene-derived artificial small RNAs (sRNAs) can induce gene silencing in some insect pests, nematodes, fungi and oomycetes, a phenomenon called host-induced gene silencing (HIGS) (Jahan et al., 2015; Nowara et al., 2010). However, the limitation associated with HIGS is the requirement for the generation of transgenic crop plants, which is a significant concern to consumers and its public acceptance is problematic in many countries. Moreover, HIGS is restricted to plants with established transformation methods, thus limiting the number of crop plants where this strategy can be applied. However, this limitation can be overcome by exogenous application of dsRNAs or sRNAs targeting pathogen genes essential for disease development. Recent studies have shown that spraying dsRNAs and sRNAs that target essential pathogen genes on plant surfaces can confer efficient and sustainable crop protection (Cai et al., 2018; Koch et al., 2016; Weiberg et al., 2013). Also called spray-induced gene silencing (SIGS), this strategy of disease control is more environmentally friendly as it leaves no chemical residues in crops and inhibits only the target organisms due to sequence specificity.

This study aims to evaluate the potential for RNAi-based spray technologies to control late blight disease in a sustainable and environmentally benign way. As a first step to ascertain whether *P. infestans* sporangia can take up dsRNA directly from their surroundings, we treated sporangia of *P. infestans* expressing Green Fluorescent Protein (GFP-*P. infestans*) with *in vitro* synthesized dsRNAs homologous to GFP (dsRNA^*GFP*^). A 436 bp dsRNA fragment derived from the *GFP* gene was labelled by incorporating Cyanine 3-UTP (Enzo Life Sciences, Inc.) into *in vitro* synthesis (Cy3-dsRNA^*GFP*^) using the MEGAscript RNAi Kit (Invitrogen). After exposure to Cy3-dsRNA^*GFP*^ for 24 hours, followed by washing with nuclease-free water to remove non-specific fluorescence, GFP-*P. infestans* sporangia were imaged using an LSM 880 confocal microscope (Zeiss Microscopy GmbH, Germany). As a control, we used dsRNA synthesized using the control template provided in the MEGAscript Kit. The analysis revealed that the GFP fluorescence was significantly reduced or disappeared in the majority of sporangia compared to the control dsRNA (Cy3-dsRNA^Ct^) treatment (Fig. 1a). Furthermore, the sporangia which exhibited reduced GFP fluorescence also exhibited Cy3 fluorescence. These results suggest that the dsRNA was effectively introduced into the *P. infestans* sporangia and dsRNAs maintain their RNAi activity by silencing the target gene. We then tested if *P. infestans* can take up dsRNA sprayed on potato leaves in a detached leaf assay (DLA). Potato leaves (cultivar Bintje) were locally sprayed on defined areas on the leaves with 20 ng μL^-1^ Cy3-dsRNA^Ct^ using an automizer. Twenty-four hours post spray application, leaves were drop-inoculated with 10 µl of GFP-*P. infestans* sporangia (5×10^4^ spores ml^−1^) and incubated in a climate-controlled chamber (22 °C daytime and 20 °C night-time temperature; 16h photoperiod). Five days post-inoculation (dpi), ∼4-5 mm diameter leaf samples from the infected part of the leaf were mounted in aniline blue solution (0.1 % aniline blue in Phosphate buffered saline (PBS), pH 7) and incubated in the dark until imaged by confocal microscopy. This demonstrated co-localization of GFP and Cy3-dsRNA^Ct^ in *P. infestans* hyphae (Fig. 1b) and sporangium (Fig. 1c) indicating uptake of dsRNA^Ct^ by GFP-*P. infestans*. Aniline blue staining demonstrated that β-1,3-glucan localization in sporangia was distinct from the co-localized Cy3-dsRNA^Ct^ and GFP (Fig. 1c). Uptake of dsRNA by fungal spores from the surrounding environment, including leaves, has been previously observed in several fungal pathosystems (Koch et al., 2016; Wang et al., 2016; Weiberg et al., 2013). Oomycetes, though evolutionarily different from fungi, may act similarly in the uptake of dsRNA from the external environment. However, the exact mechanism of sRNA and dsRNA uptake is yet to be determined.

**Figure 1.**
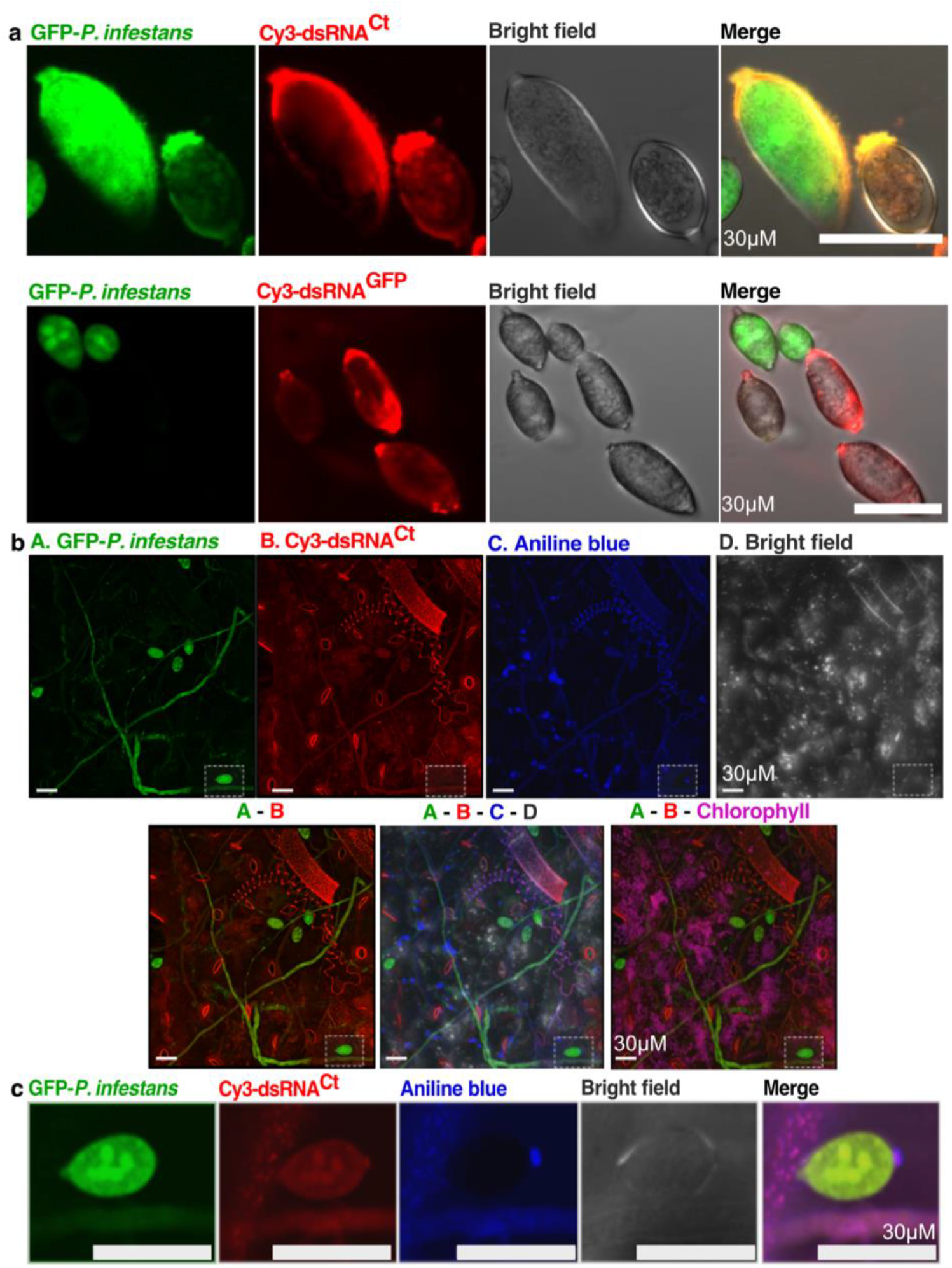
dsRNA induced gene silencing and uptake of dsRNA by *P. infestans* sporangia. **a**. Representative confocal microscopy images showing colocalization of GFP and Cy3 in dsRNA^Ct^ treated GFP-*P. infestans* sporangia (upper panel) while reduced GFP accumulation in the GFP-*P. infestans* sporangia that took up dsRNA^*GFP*^ (lower panel). Sporangia were imaged 24-hour post-treatment with dsRNA. **b**. Representative confocal microscopy images show accumulation of Cy3-dsRNA in the hyphae and sporangia of *P. infestans*, trichome, stomatal guard cells and epidermal cells of potato leaf. Images were taken with wavelengths corresponding to GFP, Cy3, and aniline blue stain. **c**. Zoomed images of the highlighted region in b, showing accumulation of Cy3-dsRNA in the sporangium. Images were taken five dpi of GFP-*P. infestans* on potato leaves.

Having established that sporangia can take up dsRNA from the surrounding environment, we next tested if the uptake of dsRNA sprayed on host potato leaves in a DLA can silence the target *P. infestans* gene. Potato leaves were locally sprayed with either 20 ng μL^-1^ dsRNA^GFP^, Cy3-dsRNA^Ct^ or water as mock treatment. One day post spraying, the leaves were drop-inoculated with GFP-*P. infestans* sporangia (10 µl of 5×10^4^ spores ml^−1^). Leaves were imaged at 5 dpi in a ChemiDoc MP imaging system (BioRad Laboratories, Inc.) using predefined settings of Alexa488 and Cy3 for visualizing GFP and Cy3 fluorescence, respectively. In line with the experiments on sporangia, the GFP fluorescence in leaves sprayed with dsRNA^GFP^ was reduced compared to Cy3-dsRNA^Ct^ and mock-treated leaves (Fig. 2a). Quantification of relative accumulation of GFP protein using immunoblot analysis with anti-GFP-HRP antibody (GF28R, Invitrogen) confirmed the observed reduction in GFP fluorescence (Fig. 2b). Real-time quantitative PCR (qRT-PCR) analysis revealed that the relative expression of *GFP* normalized to *PiActin* (NCBI P22131) was reduced by half in dsRNA^*GFP*^ compared to mock and dsRNA^Ct^ treated leaves (Fig. 2c). Notably, the relative expression of *P. infestans* actin *(PiActin)* normalized to potato actin (*StActin* NCBI XM_006345899) remained unchanged (Fig. 2d). Taken together, these results demonstrate target-specific dsRNA mediated gene silencing.

**Figure 2.**
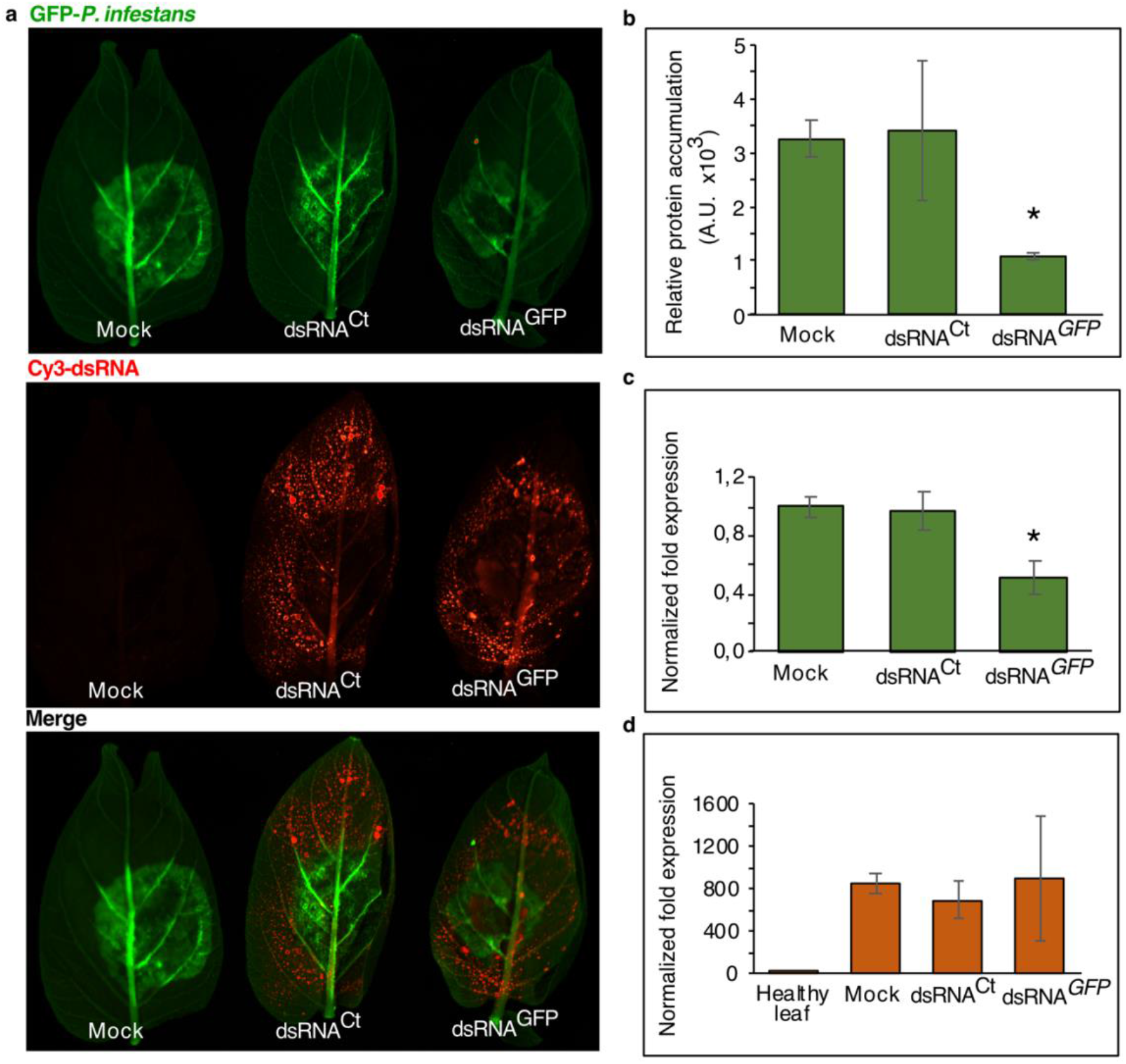
dsRNA spray mediated GFP gene silencing in *GFP-P. infestans* in detached potato leaves. **a**. Representative images showing GFP accumulation on leaves sprayed with either water (mock treatment), Cy3-dsRNA^Ct^ or Cy3-dsRNA^*GFP*^. Approximately 10 µg of dsRNA was sprayed on each leaf, left for 24 hours to dry, followed by inoculation with GFP-*P. infestans*. Leaves were imaged 5 dpi using Bio-Rad ChemiDoc at respective wavelengths for GFP and Cy3. **b**. Immunoblotting with anti-GFP-HRP antibody showing reduced GFP protein levels in dsRNA^*GFP*^ treated sample compared to mock treatment or dsRNA^Ct^. **c - d**. qRT-PCR analysis showing the downregulation of *GFP* expression in dsRNA^*GFP*^ treated samples compared to mock treatment or dsRNA^Ct^ (c). In contrast, the expression of *PiActin* is relatively unchanged (d), indicating the specificity of dsRNA^*GFP*^ mediated silencing. Asterisks indicate statistically significant difference relative to mock treatment control; *P < 0.01; Student’s t-test.

To explore the potential of SIGS as a tool to control potato late blight disease, we targeted a variety of *P. infestans* genes reported to be essential for pathogenesis, expressed at different stages of the infection cycle, and an agrochemical target. These genes included, guanine-nucleotide binding (G) protein β-subunit (*PiGPB1*; PITG_06376; XP_002998508), oxysterol binding protein (*PiOSBP*; PITG_10462; XP_002902250), haustorial membrane protein (*PiHmp1*; PITG_00375; XP_002908980), cutinase (*PiCut3*; PITG_12361; XM_002900240), and endo-1,3(4)-β-glucanase (*PiEndo3*; PITG_13567; XP_002899770).

PiGPB1 is associated with signal transduction during pathogenesis and is reported to be critical for proper sporangial development (Latijnhouwers & Govers, 2003). Through HIGS, it was previously demonstrated that targeting *PiGPB1* resulted in severe disease reduction, especially during the transition from biotrophic to the necrotrophic stage (Jahan et al., 2015). Oxysterol binding protein (PiOSBP) is the target of the oxathiapiprolin, a recent agrochemical effective against *Phytophthora* sp. (Miao et al., 2016; Miao et al., 2018; Pasteris et al., 2016). Although the exact function of OSBP is not clearly established, in other eukaryotes it is suggested to play a role in membrane-mediated lipid transport and intercellular distribution of lipid molecules (Raychaudhuri & Prinz, 2010).

Penetration and colonization of host tissue is paramount for successful infection by *P. infestans*. Penetration of the outer tissue primarily comprising cutin and β-1,4-glucans requires action of degradative enzymes including carbohydrate esterases (CE) such as cutinases and glycoside hydrolases (GH) such as endo- or exoglucanases, together known as carbohydrate-active enzymes or CAZymes (Brouwer et al., 2014; Ospina-Giraldo et al., 2010b). Here, we targeted two genes encoding such degradative proteins, *P. infestans* endo-1,3(4)-β-glucanase (PiEndo3) and *P. infestans* cutinase (PiCut3), both of which exhibit elevated transcript levels in germinating cysts (Ah-Fong et al., 2017) and thus present at the time of host tissue invasion. PiEndo3 is a GH family 81 enzyme (FungiDB; https://fungidb.org/fungidb/app/) with potential activity on cellulose and 1,3-β-glucans, both of which may be found in *P. infestans* and plants (e.g. callose in plants). PiCut3 belongs to CE family 5 (Ospina-Giraldo et al., 2010b) but the precise importance of PiCut3 in *P. infestans* pathogenicity has not been determined. The high expression of *PiCut3* during the initial stages of infection (Ospina-Giraldo et al., 2010b) suggests a role in the degradation of cutin at the outermost pathogen-plant barrier. Following penetration, the membrane-associated and infection-induced PiHmp1 protein plays a critical role in the intercellular progression and host colonization of *P. infestans* (Avrova et al., 2008). Hmp1 is considered to be necessary for haustorium formation (Avrova et al., 2008), biotrophic pathogen structures which extend into host cells for delivery of defence suppressing effector proteins (Boevink et al., 2011; Kagda et al., 2020).

To investigate the effect of targeted dsRNA treatments on development of *P. infestans*, detached potato leaves were sprayed with 500 µl of 20 ng μL^-1^ dsRNA^Ct/GFP^ as controls, or dsRNA specific to the individual target genes outlined above. At 24h post spraying the leaves were drop-inoculated (10 µl of 5×10^4^ spores ml^−1^) of *P. infestans* isolate 88069. At 5 dpi, trypan blue staining of the inoculated leaves was carried out to determine the progression of *P. infestans*. Briefly, leaves were incubated in the trypan blue staining solution (Koch & Slusarenko, 1990) for 30 min, followed by a single wash with 100 % ethanol and overnight incubation in 100 % ethanol at room temperature (Fernández-Bautista et al., 2016). Leaves were then carefully transferred to a 20 % glycerol solution and were imaged using a scanner (Epson V850Pro). Although normal disease progression was observed in the mock and dsRNA^Ct/GFP^ treated leaves, *P. infestans* development was severely inhibited in the dsRNA^*PiGPB1*^, dsRNA^*PiEndo3*^, dsRNA^*PiCut3*^ and dsRNA^*PiHmp1*^ treated leaves (Fig. 3a) but not in the dsRNA^*PiOSBP*^ sprayed leaves. Quantification of the area of infection sites using ImageJ revealed that the mean area of infection sites in the mock and control were 2.2 and 2.4 cm^2^, respectively. The mean area of infection sites in the dsRNA treated samples ranged from 0.6 to 1.24 cm^2^ indicating an apparent reduction in the area of infection in the DLAs (Fig. 3b). As no significant reduction in *P. infestans* growth was observed in the dsRNA^*PiOSBP*^ treated leaves (mean area of infection 2.69 cm^2^), the DLAs were limited to two replications for dsRNA^*PiOSBP*^ treatment, while DLAs for each of the other targets were repeated at least five times with six leaves in each experiment (n = ∼30; n = 15 for dsRNA^*PiOSBP*^). To confirm that the observed inhibition of disease progression was due to dsRNA mediated gene silencing, the relative gene expression of the target genes in the infected dsRNA treated leaves was quantified using qRT-PCR. Total RNA was extracted from 5 dpi dsRNA treated leaves (Qiagen RNeasy Plant total RNA extraction kit) followed by DNase treatment (Turbo DNA-free kit, Ambion) and cDNA synthesis (qScript SuperMix, Quantabio). Undiluted cDNA (1 µL) was used as a template for qRT-PCR (DyNAmo Flash SYBR Green kit, Thermo Scientific). The target gene transcript levels were normalized to the expression of reference gene *PiActin* (Vetukuri et al., 2011). Compared to the dsRNA^Ct^ treated samples, a 2.5-, 1-, 1.5-, and a 2-fold decrease was observed in *PiGPB1, PiHmp, PiCut3* and *PiEndo3* transcript levels in each of the respective treatments, suggesting that the observed reduction in the *P. infestans* disease progression is indeed a result of dsRNA mediated targeted gene silencing (Fig. 3c).

**Figure 3.**
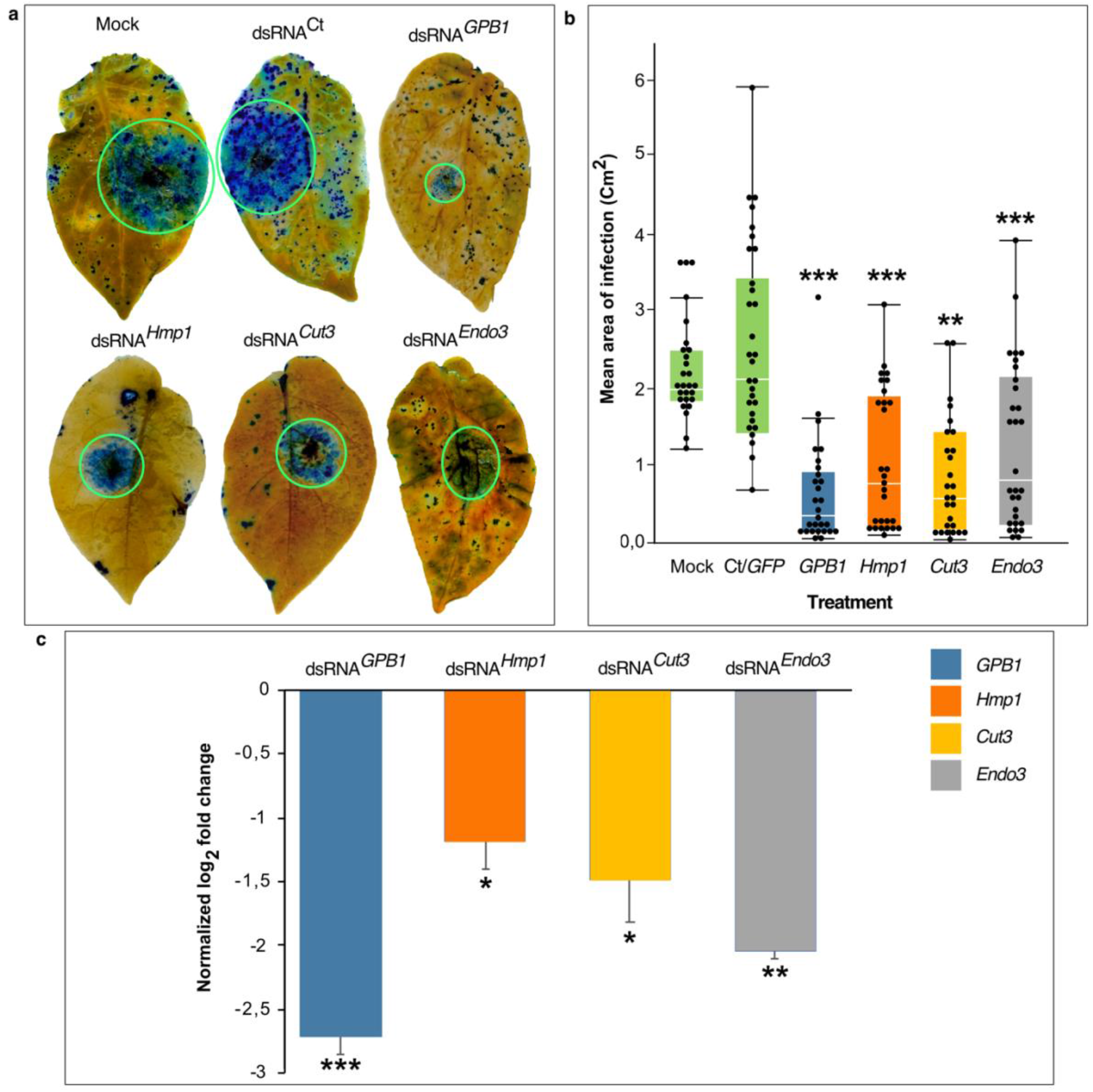
Effect of SIGS on *P. infestans* disease progression. **a**. Representative pictures of trypan blue stained potato leaves showing *P. infestans* 88069 disease progression at 5 dpi on leaves sprayed with water (mock treatment), dsRNA^Ct^, dsRNA^*PiGPB1*^, dsRNA^*PiEndo3*^, dsRNA^*PiCut3*^ or dsRNA^*PiHmp1*^. **b**. Box plot showing the quantification of area of disease progression for each dsRNA sprayed (n = 30). **c**. qRT-PCR analysis showing the relative gene expression of each target upon treatment with respective dsRNA. The Cq values of target genes were normalized to the Cq values of *PiActin*. ***P < 0.0001, **P < 0.001, *P < 0.01; Control, Dunnet’s test.

Prior studies that have noted that the formation of sporangia during 36-48 hours post-infection is critical for the disease progression and transition from biotrophic to necrotrophic phase (Judelson & Blanco, 2005). To investigate if SIGS mediated inhibition of disease progression was also associated with defects in sporulation, we carried out a microscopic examination of disease lesions (Leica MDG41 stereo microscope). Our analysis revealed that treatment with dsRNA^*PiGPB1*^ resulted in severe sporulation inhibition (Fig. 4a-b, f) compared to the dsRNA^Ct^ treatment. This agrees with Jahan et al. (2015) and Latijnhouwers & Govers (2003) who demonstrated the formation of fewer and deformed sporangia through silencing in hp-PiGPB1 transgenic plants and transcriptional silencing in *P. infestans* stable transformants, respectively. Also, in agreement with those studies, treatment with dsRNA^*PiGPB1*^ did not appear to disrupt the germination of the sporangia or mycelial progression, as mycelia could be seen emerging from stomata (Fig. 4b).

**Figure 4:**
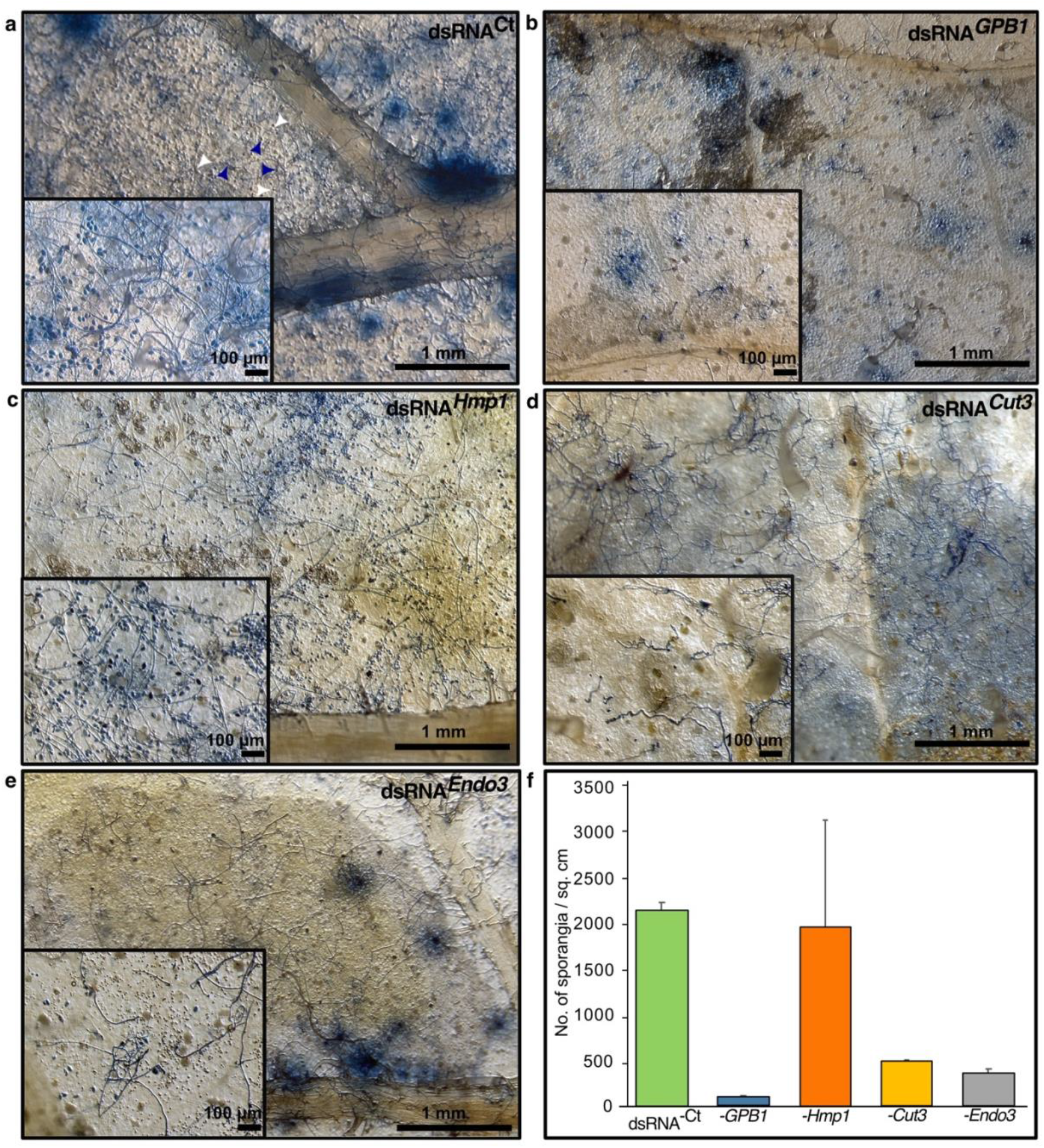
Effect of SIGS on *P. infestans* morphology. **a-e**. Representative stereo microscope images showing the effect of respective dsRNAs on the *P. infestans* morphology on detached potato leaves. Leaves were stained with trypan blue followed by imaging on a Leica stereomicroscope with 3.2x objective. Images in the inset were taken with 12x objective. **f**. Average number of sporangia per cm^2^ of infection. Sporangia were counted manually from the images taken using the 12x objective. The graph represents mean sporangia count from three images taken from individual leaves.

As expected, silencing genes encoding CAZymes and Hmp1 did not result in severe inhibition of sporulation (Fig. 4c). However, the number of sporangia observed was significantly lower in the leaves treated with dsRNA^*PiEndo3*^ and dsRNA^*PiCut3*^ (Fig. 4d - f) suggesting that the lower sporangial count could be as a result of reduced disease progression rather than a direct effect on the sporangial development.

*P. infestans* cell walls contain both cellulose and 1,3-β-glucan, which may be substrates for PiEndo3, potentially leading to remodeling of the wall. However, our findings do not indicate any direct effect of *PiEndo3* on the growth and development of *P. infestans*, thus it is likely that this enzyme acts on plant polysaccharides, and decreased disease progression can be attributed to the silencing of *PiEndo3* expression. Further analysis focused on more detailed phenotyping might be valuable in unravelling the functions of this and other *P. infestans* glucanases. Interestingly, treatment with dsRNA^*PiCut3*^ resulted in a smaller and aberrant mycelial phenotype (Fig. 4d, inset), probably owing to erratic penetration, causing nutrient starvation. This is consistent with the expected role of cutinase in facilitating host penetration by *Phytophthora* species by breaking down cutin and enabling plant cell wall disintegration (Ospina-Giraldo et al., 2010a; Zerillo et al., 2013).

Although disease progression was significantly lower in the dsRNA^*PiHmp1*^ treated leaves, the number of sporangia per cm^2^ was not reduced (Fig. 4c, f). These results corroborate Avrova et al. (2008) who showed that dsRNA-mediated transient silencing of *PiHmp1* produced a similar number of sporangia compared to the control when grown in agar culture, even though the infection was suppressed compared to the control non-homologous dsRNA-treated lines.

Our observations show that not all sporangia take up the dsRNA to levels detectable using conjugated Cy3 dye. Hence, optimizing formulation of dsRNA to facilitate dsRNA uptake will be crucial to successfully use SIGS for disease control (San Miguel & Scott, 2016; Yan et al., 2020). It is also possible that spray-applied dsRNAs can enter *P. infestans* as spores germinate on the leaf surface; this is evidenced here (Fig. 1b, c) where Cy3 labelled dsRNA can be seen labelling pathogen hyphae. A further question arises regards the nature of the dsRNAs taken up by *P. infestans*. That is, it remains to be determined if *P. infestans* takes up the long dsRNA molecules, or whether the gene silencing is indirect, with dsRNAs first entering plant cells where they are processed into siRNAs prior to entry into *P. infestans*. However, it has been shown that long dsRNAs can enter *P. infestans* protoplasts derived from hyphae (Whisson et al., 2005), so at least some of the spray-applied dsRNAs are likely to enter *P. infestans* directly.

Although dsRNA mediated SIGS has been reported in other pathogens (Cai et al., 2018; Koch et al., 2016; Weiberg et al., 2013), it is yet to be proven if this gene silencing is as a result of plant RNAi machinery or the pathogen RNAi mechanism. Our analysis using CLSM showed uptake of Cy3-dsRNA in the *P. infestans* mycelium and sporangia indicates that the site of dsRNA processing for RNA silencing is located in the pathogen (Fig. 1b, c). Further analysis including sRNA sequencing from the dsRNA sprayed leaves, could prove if the plant RNA silencing machinery is processing the sprayed dsRNAs. Our findings while preliminary, provide a proof of concept for SIGS applications to control potato late blight and other *Phytophthora* diseases. This also has implications for the development of potential alternative reverse genetic tools in challenging organisms like *P. infestans*.

## ACKNOWLEDGEMENTS

This work has been supported by FORMAS (2019-01316), The Swedish Research Council (2019-04270), NKJ-SNS - Dialogue Biocontrol network (NKJ-SNS 06), Carl Tryggers Stiftelse för Vetenskaplig Forskning (CTS 20:464), The Crafoord foundation (20200818), Partnerskap Alnarp (1317/Trg,VO/2020) and Alnarp stipendiekommitténs. MD was supported by FORMAS (2018-01420). SCW acknowledges financial support from the Scottish Government Rural and Environment Science and Analytical Services Division (RESAS). The authors have no conflicts of interest to declare.

## DATA AVAILABILITY STATEMENT

All gene sequences used in this study have been obtained from NCBI GenBank; accession numbers are given in the text of the article. Primers used in this study are available as supplementary material.

