## Supplementary data for "Spray-induced gene silencing as a potential tool to control potato late blight disease"

Table S1. List of primers used in the study

| **Primers used for dsRNA synthesis** | **Primer sequence (5'-3')** (T7 sequence) |
|---|---|
| T7 EGFP dsRNA.FOR1 | GTAATACGACTCACTATAGGGGACGTAAACGGCCACAAGTT |
| T7 EGFP dsRNA.REV2 | GTAATACGACTCACTATAGGGAGTTCACCTTGATGCCGTTC |
| T7 PiGPB1 dsRNA.FOR1 | GTAATACGACTCACTATAGGGATGTTTATTTCGGGCTCGTGTGA |
| T7 PiGPB1 dsRNA.REV1 | GTAATACGACTCACTATAGGGTAGATATGCGCTCCGGAAGT |
| T7 OSBP dsRNA.FOR 1 | GTAATACGACTCACTATAGGGGAACCTCGAACCAATCTGGA |
| T7 OSBP dsRNA.REV 1 | GTAATACGACTCACTATAGGGGTTAAGCATGGCGTTGGATT |
| T7 PITG_00375 dsRNA.FOR1 | GTAATACGACTCACTATAGGTTGTTGACCAGCTCGTTGAG |
| T7 PITG_00375 dsRNA.REV1 | GTAATACGACTCACTATAGGCCATCACCCTTCTTCTTCCA |
| T7 PITG_12361 dsRNA.FOR1 | GTAATACGACTCACTATAGGCAACCACGTCGTGTCTATCG |
| T7 PITG_12361 dsRNA.REV1 | GTAATACGACTCACTATAGGGTTGCAGAACTCAATGGCCT |
| T7 PITG_13567 dsRNA.FOR1 | GTAATACGACTCACTATAGGCGTCGAGGATCTAGGCAGTC |
| T7 PITG_13567 dsRNA.REV1 | GTAATACGACTCACTATAGGGACGAGAAGCAGCGATAACC |
| **qPCR Primers** | **Primer sequence (5'-3')** |
| EGFP qPCR.FOR | GACCACTACCAGCAGAACACC |
| EGFP qPCR.REV | CGCTTCTCGTTGGGGTCTTT |
| PiGPB1 qPCR.FOR | TTCCGGAGCGCATATCTACC |
| PiGPB1 qPCR.REV | TCTTGAGTAGCGTGTCCCAG |
| qPCR_00375.FOR | GTGAAGGTGGTACCGAAGGA |
| qPCR_00375.REV | TCTTGCCCTTGTTCTTGTCC |
| qPCR_12361.FOR | CCGACGTGCATGTTGTCTTC |
| qPCR_12361.REV | GGTTCGAGGTGATCCCACTG |
| qPCR_13567.FOR | AACTATCGCAAGGGCAGGTT |
| qPCR_13567.REV | CGTCCACCGATGTCTGAGTT |
